# AN OPTIMIZED AND DRUGGABLE HUMAN KERATINOCYTE AND IPSC-DERIVED SENSORY NEURON CO-CULTURE SYSTEM FOR ATOPIC ITCH

**DOI:** 10.64898/2026.05.09.724000

**Authors:** Hendrik Mießner, Benjamin Al, Hendrik Reuter, Judith Anna Seidel, Ewan St. John Smith

**Affiliations:** Department of Dermatology, Beiersdorf AG; Hamburg, Germany; Department of Pharmacology, University of Cambridge; Cambridge, United Kingdom

**Author notes:** Corresponding author. Name: Ewan St. John Smith.

## Abstract

Atopic dermatitis (AD) is a highly prevalent, relapse-remitting, inflammatory skin disease, the hallmark symptom of which is chronic itch. Mechanisms underlying AD itch are multifactorial, involving various cells, receptors, and mediators. Developing a physiologically relevant, human model system for AD itch research and drug development is crucial. To this end, human induced pluripotent stem cell-derived sensory neurons (iPSCSNs) were cultured with human primary keratinocytes to form deconstructed skin models. Using Ca^2+^-imaging in a direct contact, 2.5D co-culturing format, which mimics natural skin innervation and permits both paracrine exchange and juxtacrine signaling, iPSCSNs exhibited functional TRPA1 responses not seen in monotypic iPSCSN cultures or in iPSCSNs conditioned with keratinocyte medium. Different AD-associated cytokines were used to stimulate the co-culture systems to mimic an inflamed lesional skin environment, whereby TNF was found to increase iPSCSN chemosensitivity. Finally, both TRPA1 and JAK1/2 inhibition reduced iPSCSN responses to pruritogens (TSLP, IL-31), thus supporting TRPA1 as a therapeutic target for AD itch in humans. This study demonstrates that human deconstructed skin models can be a useful tool in AD and broader pruritus research.

**GRAPHICAL ABSTRACT:** 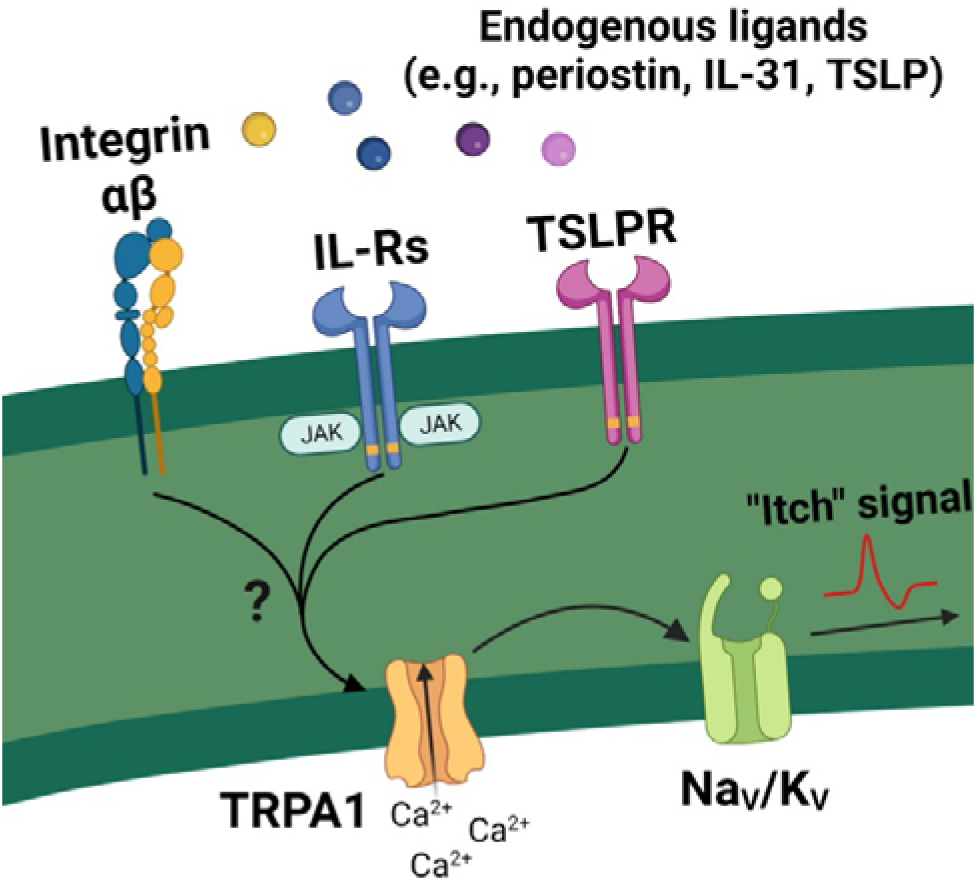

## INTRODUCTION

The skin is densely innervated by an array of specialized sensory neurons, some subserving touch, others translating nociception and the sensation of pain, and others mediating pruriception and the sensation of itch (Ständer and Schmelz, 2024). In chronic skin diseases such as atopic dermatitis (AD), which is characterized by remitting inflammatory lesions, itch is a major symptom (Silverberg et al., 2018, Ring, 2021). Chronic itch causes excessive scratching, which disrupts the skin barrier and exacerbates inflammatory responses (Yosipovitch et al., 2019), as well as impacting sleep and the psychological wellbeing of those affected (Mann et al., 2020, Weisshaar, 2021). Although recent years have seen new treatments enter clinical trials (Mahmoud et al., 2024), greater mechanistic understanding of disease processes is required to further develop therapies for chronic itch. For this, physiologically relevant experimental models are needed.

Advancements in disease models include numerous mouse models of pruritus (Donglang et al., 2021). Some screenable *in vitro* models of mostly human origin have emerged as promising alternatives (Mießner et al., 2022). However, many *in vitro* itch model systems focus solely on sensory neurons. To more realistically mimic neuron-skin interactions, co-culture models with sensory neurons and keratinocytes have been established (Roggenkamp et al., 2012, Tsantoulas et al., 2013), which is incredibly important considering the growing understanding of neuron – non-neuron interactions in the processing of sensory signals (Lin et al., 2024, Smith et al., 2025). In AD, scratching-induced epidermal secretion of nerve growth factor (NGF) and glial cell line-derived neurotrophic factor (GDNF) is thought to contribute to increased sensory neuron activity and innervation (Roggenkamp et al., 2013, Tominaga and Takamori, 2014). A crucial role for keratinocyte-mediated neuronal regulation is further supported by studies showing that conditioned keratinocyte medium enhances human induced pluripotent stem cell-derived sensory neuron (iPSCSN) substance P release (Guimaraes et al., 2018) and that primary keratinocytes attract iPSCSN neurites (Belamadni et al., 2022). Measuring pruriception includes assessing the roles of ion channels associated with itch, such as transient receptor potential vanilloid-type 1 (TRPV1) and ankyrin-type 1 (TRPA1) (Wilson et al., 2013a, Kittaka and Tominaga, 2017). Additionally, measuring responses to AD pruritogens such as keratinocyte-derived thymic stromal lymphopoietin (TSLP) (Berna et al., 2021) and Th2-associated interleukin-31 (IL-31) (Meng et al., 2018, Silverberg et al., 2021, Jha et al., 2025) is essential as both directly activate sensory neurons (Wilson et al., 2013b, Cevikbas et al., 2014).

In this study, we introduce a human deconstructed skin model using iPSCSNs and keratinocytes that simulates AD and is accessible for pharmacological testing. Co-culture reveals that keratinocyte contact/innervation, and the AD-associated cytokine tumor necrosis factor (TNF) increase iPSCSN pruritogen sensitivity. Using this human model system, we demonstrate that TRPA1 inhibition can reduce TSLP-/IL-31-induced neuronal activity.

## RESULTS

### Section 1: iPSCSNs display nociceptive neuron-like gene expression and electrophysiological properties after long-term culture

We cultured iPSCSN progenitors (Axol Bioscience) for 8 weeks until they reached a nociceptive neuron-like maturation state (Figure 1a), when morphology and size (20-50 µm diameter) resembled primary dorsal root ganglia (DRG)-derived sensory neurons (Anand et al., 2006) with a dense neurite network (Figure 1b). Like primary human DRG and nerve fibers innervating human skin, 8-week iPSCSNs expressed either neurofilament-H (NF-H), peripherin or both (Figure 1c). Confirming successful differentiation to peripheral sensory neurons, decreased expression of the stem cell/progenitor markers *NEUROG1*, *OTX2* and *SOX10* was observed at 8-weeks vs. t0 (Table 1, Figure S1a) alongside increased gene expression for peripherin (*PRPH)* and the neurotransmitter substance P (*TAC1*); substance P was also detected using immunohistochemistry (Figure S1b; CGRP could not be detected, data not shown). Like mature mammalian sensory neurons, 8-week iPSCSNs also displayed significantly increased expression of the voltage-gated sodium channel subunits Na_V_1.7 (*SCN9A*) and Na_V_1.8 (*SCN10A*) (Figure S1c). Given their role in pruriception, TRP channel expression was also examined. Gene expression analysis detected *TRPV1* in iPSCSN progenitors at t0 (based on raw data, Ct-value <30) with no increase in expression following 8-week differentiation, whereas expression of *TRPM8* and *TRPA1* decreased and increased respectively (Figure S1d); immunocytochemistry corroborated these results (Figure S1e). In addition, the P2X3 receptor (*P2RX3*), an ATP-gated ion channel implicated in nociception, was detectable in ∼40% of iPSCSNs at both t0 and following 8-week differentiation (Figure S1d/e). Adding potential utility for AD/itch-associated research, TSLPR (thymic stromal lymphopoietin receptor, *CRLF2*) showed much greater expression after 8 weeks (Figure S1f).

**Figure 1:**
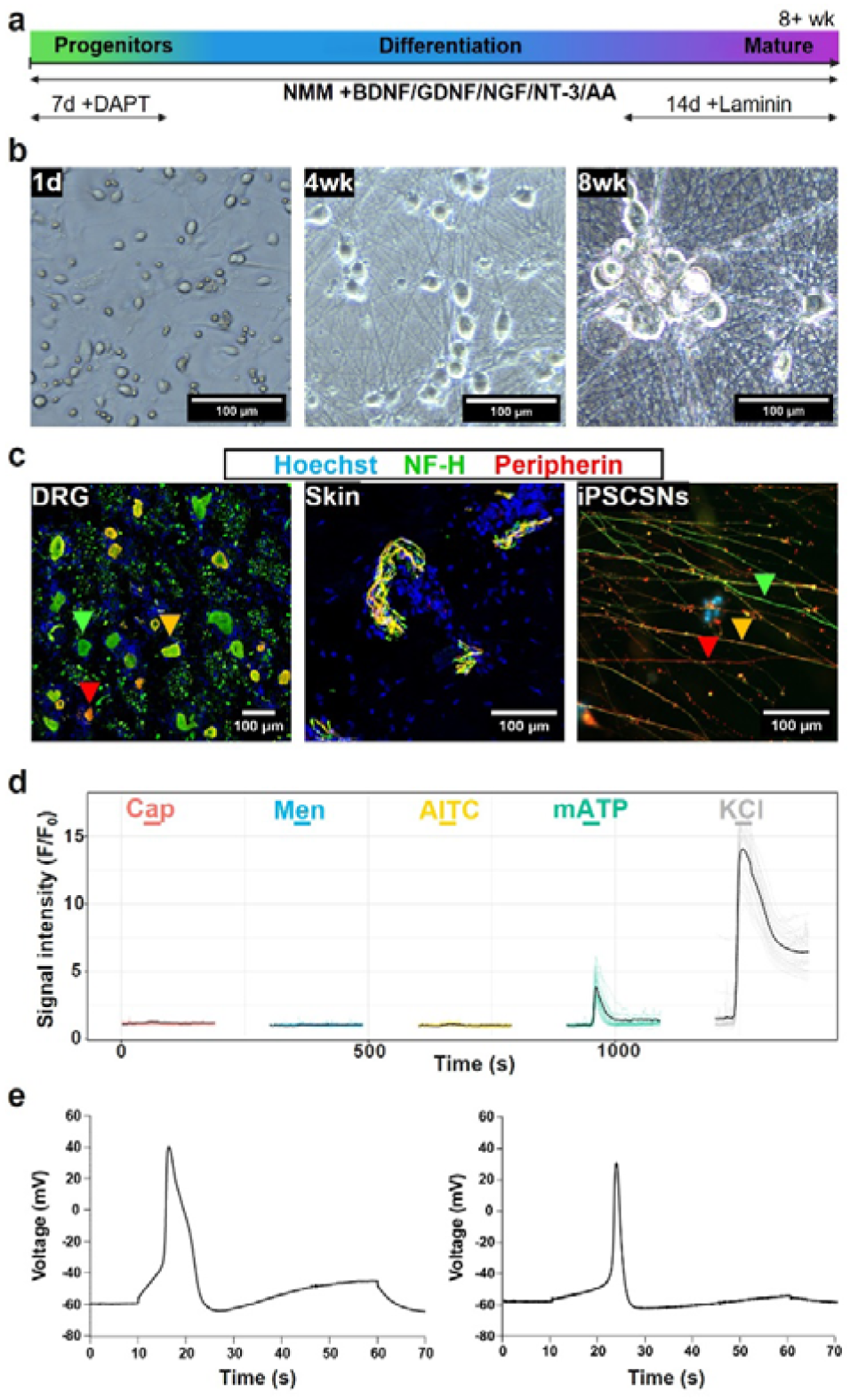
Morphology and functional characterization of iPSCSNs after long-term culture. (a) 8-week long treatment and culture timeline from progenitor state to mature sensory neurons. (b) iPSCSNs after 1 day, 4 and 8 weeks of culture. (c) Immunohistochemistry showing the expression of neurofilament-H (NF-H), peripherin (PRPH) and nuclei (Hoechst) in human dorsal root ganglia (DRG), skin and iPSCSN. Arrowheads indicate neurons/neurites expressing NF-H and PRPH exclusively or combined. (d) Example Ca^2+^-imaging recording of positively responding iPSCSNs consecutively treated for 20s with capsaicin (1 µM), menthol (250 µM), allyl isothiocyanate (AITC, 100 µM), α,β-methylene ATP (mATP, 100 µM), KCl (50 mM). (e) Patch-clamp recording showing a nociceptive neuron-like AP at rheobase threshold with its characteristic inflection in the falling phase (left) compared to a narrow AP without inflection (right). Scale bars represent 100 µm. See also Figure S1 and S2 for complementary data. NMM, Neuronal maturation/maintenance medium; BDNF, brain-derived neurotrophic factor; GDNF, glial-derived neurotrophic factor, NGF, nerve growth factor; NT-3, Neurotrophin-3; DAPT, tert-Butyl (S)-{(2S)-2-[2-(3,5-difluorophenyl)acetamido]propanamido}phenylacetate; Cap, Capsaicin; Men, Menthol; AITC, Allyl isothiocyanate; mATP, α,β-methylene ATP; KCl, potassium chloride.

**Table 1:**
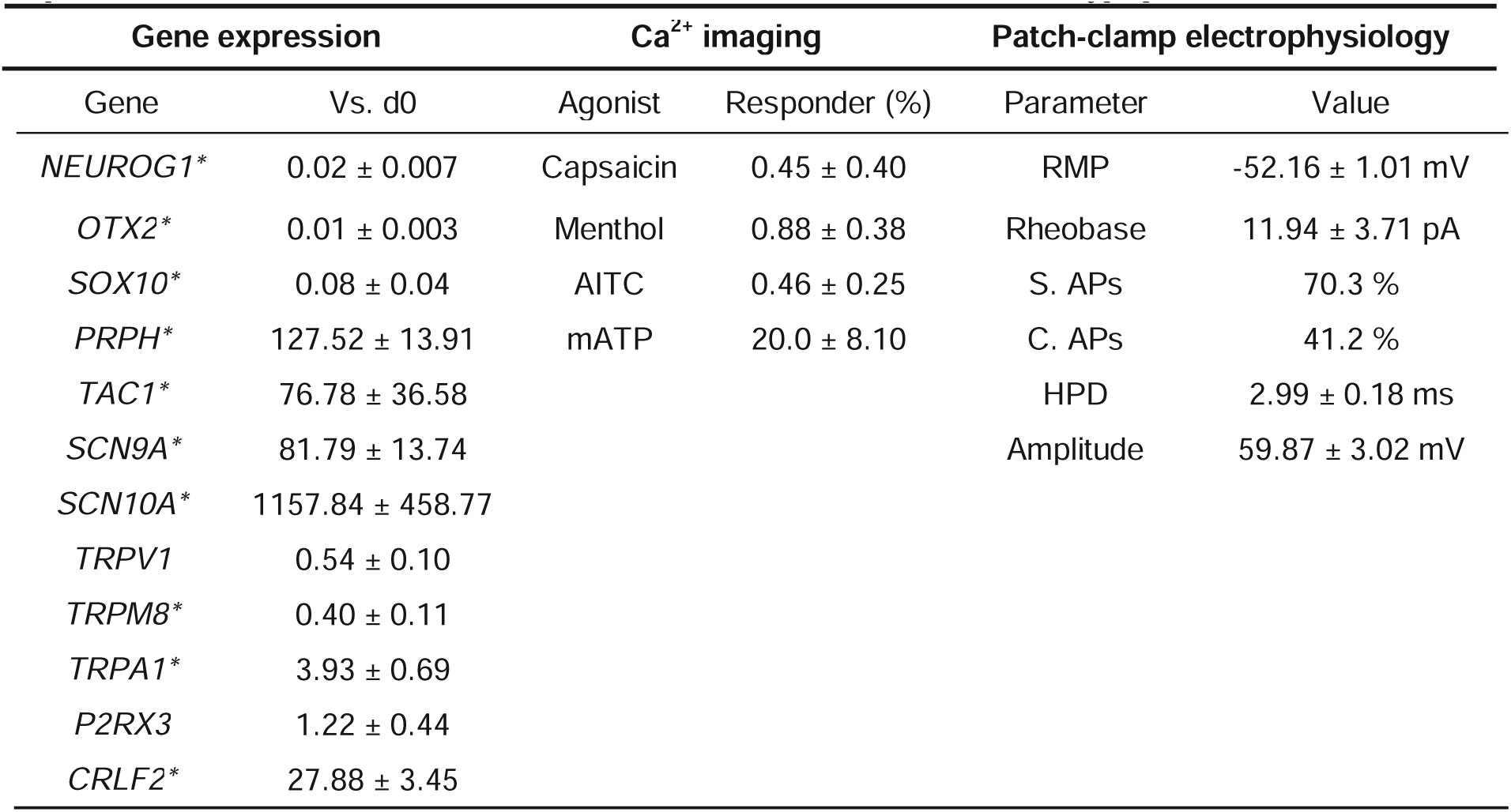
Gene expression, Ca^2+^-imaging and patch-clamp electrophysiology parameters of 8-week differentiated iPSCSNs for nociceptive neuron-like characterization. Asterisk* = significant difference between 8-week and d0 gene expression. See Figures S1/2 for further information. RMP, Resting membrane potential; S. APs, Spontaneous APs; C. APs, Consecutive APs; HPD, Half-Peak Duration; AHP, Afterhyperpolarization.

Taken together, 8-week-differentiated iPSCSNs exhibit gene expression characteristic of a mature, nociceptive neuron-like phenotype.

For functional assessment of 8-week iPSCSNs, cells were loaded with the non-ratiometric Ca^2+^-indicator Calbryte-520 and tested for responses to agonists of TRPV1 (capsaicin, Cap), TRPM8 (menthol, Men), TRPA1 (allyl isothiocyanate, AITC) and P2RX3 (α,β-methylene ATP, mATP) (Figure 1d). Of cells responding to KCl, 20.0 ± 8.1% also responded to mATP, whereas responses rarely crossed the activation threshold for Cap, Men and AITC (<1% on average, Table 1, Figure S1g). The lack of response to Cap, Men and AITC contrasts with mRNA and protein (Figure S1d/e) expression data, but was consistent across 5 batches of differentiation, 11 experiments and 1834 cells in total.

Action potential electrogenesis in iPSCSNs was next investigated using patch-clamp electrophysiology. When entering whole-cell configuration, the resting membrane potential was-52.16 ± 1.01 mV (Table 1, Figure S2a) with 70.3% of cells exhibiting spontaneous firing (Figure S2b/c), which mostly stopped shortly after cell-access or following manual current injection. Scoring such neurons as having an action potential threshold of 0 pA, the iPSCSN rheobase was 11.94 ± 3.71 pA; of those iPSCSNs that did not spontaneously fire, the rheobase was 39.09 ± 7.19 pA. Examining further action potential parameters, in 7 dishes/biological replicates, 0-20 % featured an inflection during repolarization, which is a feature of nociceptive neurons (Lawson et al., 1996). The remaining iPSCSNs displayed narrow action potentials with no inflection during repolarization (Figure 1e). Further action potential parameters were as follows: half-peak duration (2.99 ± 0.18 ms), afterhyperpolarization time (34.94 ± 2.75 ms) and amplitude (59.87 ± 3.02 mV) (Figure S2a). When currents above rheobase were injected for 500 ms, 55% of iPSCSNs fired multiple action potentials (Figure S2b). In summary, whilst exhibiting some key aspects of sensory neuron function, e.g. action potential electrogenesis and P2X3R-mediated currents, iPSCSNs lacked responses to certain TRP channel agonists, contrasting with what expression data would predict.

### Section 2: Contact with keratinocytes improves iPSCSN differentiation and chemosensitivity

To simulate the skin environment, where sensory nerve endings interact with surrounding keratinocytes and mediators they release, we created two iPSCSN / primary human epidermal keratinocyte (HEK) co-culture systems. First, an insert-based system with shared culture medium but no direct contact was tested, a system enabling HEK metabolites to pass through the membrane they were cultured on to iPSCSNs growing in the chamber below (medSN, Figure 2a). To support modulation of early-stage neuron development for as long as possible, whilst also considering the limited ability of HEK to survive prolonged periods in culture, medSN differentiation was limited to 5 weeks (2 weeks without keratinocytes, 3 weeks co-culture). After 5-week medSN co-culture, results revealed similar responses to those observed in monoculture, i.e. no robust responses to TRP channel agonists (Figure 2b), but responses were observed in occasional experiments probing for mATP reaction (data not shown).

**Figure 2:**
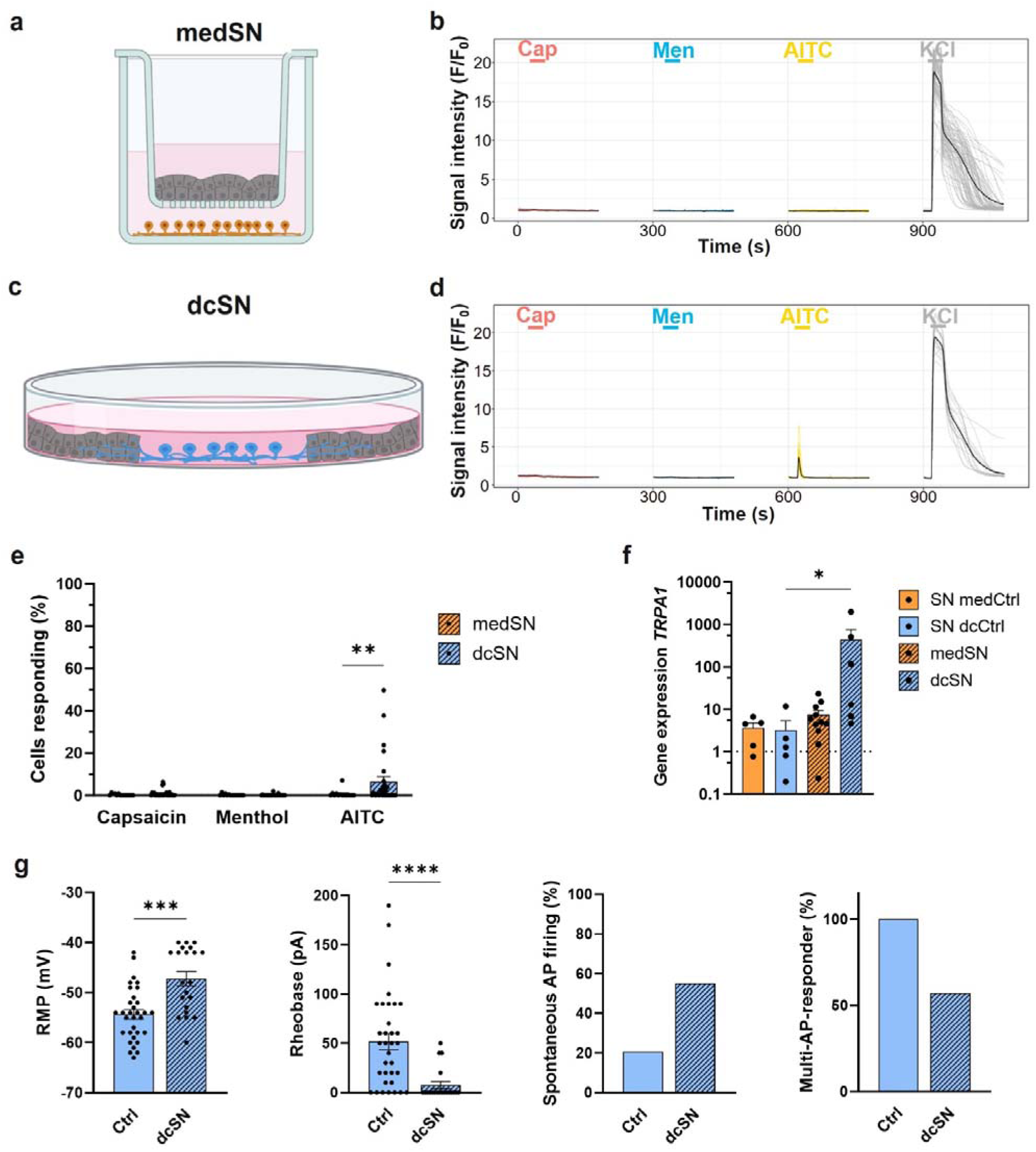
Effect of keratinocytes on iPSCSNs in different co-culture systems. (a) Primary human epidermal keratinocytes (HEK) grown in 24-well plate inserts and added to d14 iPSCSN culture (medSN, medCtrl without HEK). (b) Example Ca^2+^ responses of iPSCSNs in medSN consecutively treated for 20s with capsaicin (Cap, 1 µM), menthol (Men, 250 µM), AITC (100 µM), KCl (50 mM) and 5 min washes in between). (c) After 14 days of iPSCSN culture in the middle of 35mm dishes, primary HEKs were added to the outer perimeter for co-culture with direct contact (dcSN, dcCtrl without HEK) and (d) representative Ca^2+^ imaging (AITC-positive traces). (e) Percentage of cells reacting to aforementioned stimuli (KCl-positive-only; Two-way ANOVA with Tukey’s multiple comparisons test, n≥8 individual cultures). (f) TRPA1 gene expression changes (normalized to iPSCSN t0, relative fold-change, 2^-ΔΔCt^, Kruskal-Wallis test with Dunn’s multiple comparisons test, n≥5 individual cultures). (g) Comparison of analyzed Ctrl and dcSN cells using patch-clamp electrophysiology with regards to RMP (resting membrane potential) and rheobase, percentage of spontaneous and multi-action potential firing cells (Mann-Whitney-test, n≥32/20 (Ctrl/dcSN) cells from 5/4 individual cultures). Data are presented as mean ± SEM. See also Figure S3 for complimentary data. (*=p<0.05; **=p<0.01; ***=p<0.001; ****=p<0.0001)

In a second iPSCSN/HEK co-culture system, iPSCSNs were seeded in the middle of a 35 mm dish and cultured for 2 weeks before adding HEKs to the perimeter of the dish, thus enabling both direct contact and mediator exchange (dcSN, Figure 2c). The dcSN approach led to a boundary layer where iPSCSN neurites innervated HEK and appeared to stop further HEK expansion toward iPSCSNs. Contact inhibition initiated HEK differentiation (Charest et al., 2009) and thus growth in height, which produced cell layer bulges and a 2.5-dimensional culture (Figure S3a). Ca^2+^-imaging of dcSN revealed iPSCSN responses to AITC application (Figure 2d) at a significantly greater frequency than was observed in medSN cultures (Figure 2e). Additionally, TRPA1 gene expression was significantly increased in dcSN (but not medSN) compared to the control culture without HEK (dcCtrl/medCtrl, Figure 2f), which likely underpins the increased AITC-sensitivity observed. Regarding electrophysiological parameters compared to dcCtrl neurons, dcSNs exhibited a more depolarized resting membrane potential (-54.34 ± 0.94 mV vs.-47.24 ± 1.41 mV), as well as a lower rheobase (51.47 ± 8.28 pA vs. 7.5 ± 3.62 pA). In addition, a greater proportion of dcSNs exhibited spontaneous firing (26.47% vs. 80.95%), although fewer dcSNs fired multiple action potentials in response to 500 ms stimulation (100% vs. 61.9 %, Figure 3g). Further action potential parameters were as follows (dcCtrl vs. dcSN): half-peak duration was longer in dcSNs (1.45 ± 0.11 vs. 2.20 ± 0.14 ms), afterhyperpolarization time similar (22.40 ± 2.67 vs. 21.32 ± 2.81 ms) and amplitude smaller (86.19 ± 3.57 vs. 62.22 ± 3.52 mV). In summary, culturing iPSCSNs in direct contact with HEKs created a deconstructed skin model that changed the expression profile, chemosensitivity, and excitability of iPSCSNs, enabling this paradigm to be used for investigating the role of TRPA1 while decreasing the iPSCSN maturation time to 5 weeks.

**Figure 3:**
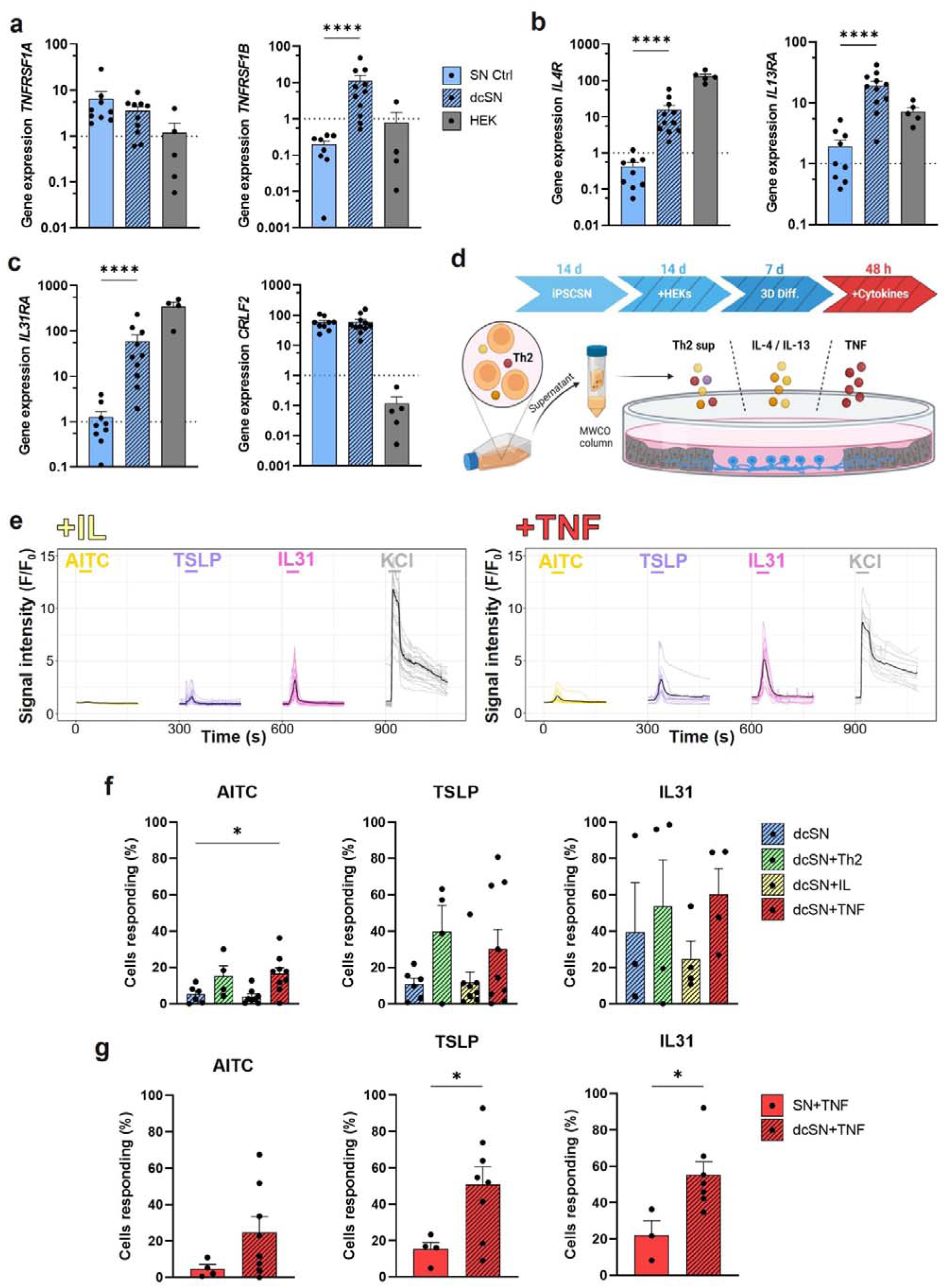
Modulation of iPSCSN pruritogen-responsiveness through co-culture with AD-associated cytokines. (a) Gene expression of *TNFRSF1A* and *TNFRSF1B*, (b) *IL4R* and *IL13RA*, (c) *IL31RA* and *CRLF2* (normalized to iPSCSN t0, relative fold-change, 2^-ΔΔCt^, Mann-Whitney-test between SN Ctrl and dcSN, n≥9, n≥4 HEK for reference). (d) Schematic depiction of Th2 cell culture, supernatant collection, concentration and its addition (alternatively IL-4/-13 50 ng/mL each or TNF 20 ng/mL) to the dcSN co-culture for 48h. (e) Example Ca^2+^-imaging of dcSN +IL or +TNF stimulated for 20s with AITC (100 µM), TSLP (2 µg/mL), IL-31 (25 µg/mL), KCl (50 mM). (f) Percentage of dcSN ± AD cytokines responding to AITC, TSLP or IL-31 (KCl-positive-only, One-way ANOVA with Dunnett’s multiple comparisons test; AITC/TSLP: n≥4, IL-31: n≥3). (g) Comparison of TNF-treated SN or dcSN responding to AITC, TSLP or IL-31 (KCl-positive-only, unpaired t test, n≥3 SN+TNF, n≥7 dcSN+TNF). Data are presented as mean ± SEM, data points are biological replicates. See also Figure S3. (*=p<0.05; ****=p<0.0001)

### Section 3: TNF further increases iPSCSN chemosensitivity in an AD skin model

Continuing with the dcSN co-culture system, we next investigated how the AD-associated cytokines TNF, IL-4 and IL-13 (Danso et al., 2014), modulated iPSCSN function. Firstly, we observed that the dcSN culturing system significantly elevated iPSCSN expression of the relevant cytokine receptors *TNFRSF1B*, *IL4R* and *IL13RA* (although not *TNFRSF1A*, Figure 3a/b). Similarly, pruritogen receptors *IL31RA* and *CRLF2* (encoding TSLPR) exhibited a high level of expression for dcSNs, while *IL31RA* was not upregulated in SN control cultures (Figure 3c).

Subsequently, supernatant from activated, *in vitro* polarized Th2 cells was administered to 5-week differentiated dcSN cells for 48 h, thus mimicking the Th2 driven inflammatory cytokine environment of AD (Al et al., 2024). After seeing trends of greater chemosensitivity, although with high donor-variability, we aimed to provide a more reproducible Th2-like stimulation focusing on potential mediators. Therefore, recombinant IL-4/IL-13 (+IL, 50 ng/mL each) or TNF (+TNF, 20 ng/mL) were also tested for their ability to induce an AD-relevant phenotype in iPSCSNs (Figure 3d). IL-31 and TSLP receptors were expressed in co-culture (Figure 3c), hence the ability of AD-relevant pruritogens IL-31 and TSLP to excite iPSCSNs was measured next.

Ca^2+^ responses to TSLP (2 µg/mL) and IL-31 (25 µg/mL) were observed under all treatment conditions (Figures 3e/f). However, +IL treatment did not result in a greater number of AITC/TSLP/IL-31 responsive iPSCSNs compared to untreated dcSN (Figure 3e/f). By contrast, both +TNF and +Th2 incubation resulted in a larger proportion of cells responding to different stimuli, which was significant for AITC when treated +TNF (Figure 3e/f); responses to TSLP and IL-31 were not significantly different due to high inter-experiment variability. Comparing different conditions, TNF appears most potent at sensitizing dcSN function and therefore replaced Th2 supernatant as an AD relevant cytokine in subsequent experiments.

To verify whether pruritogen-responsiveness observed in TNF-treated iPSCSNs was dependent on the presence of HEK, the assay was repeated in the absence of HEK. TNF treatment of iPSCSNs alone (SN+TNF) resulted in only moderate numbers of cells responding to AITC, TSLP, IL-31 and the recently described pruritogen periostin (Figures 3g, S3c). By contrast, the proportion of dcSN+TNF responses to TSLP and IL-31 was significantly higher compared to SN+TNF cells, thus demonstrating the importance of HEK in creating an AD-like co-culture (Figure 3g).

### Section 4: TRPA1 inhibition reduces TSLP/IL-31-induced iPSCSN activation

In rodent models, TRPA1 has been implicated in TSLPR-and IL-31R-mediated itch signaling (Wilson et al., 2013b, Cevikbas et al., 2014). We therefore investigated whether this mechanism also occurs in human iPSCSNs using the TRPA1 antagonist A967079. We first confirmed that the inhibitor reversibly suppressed AITC-stimulated Ca^2+^ responses in dcSN+TNF (Figure 4a/b). Importantly, when applied with TSLP (Figure 4c/d) or IL-31 (Figure 4e/f), A967079 inhibited Ca^2+^ responses in a reversible manner. These results confirm a role for TRPA1 in TSLPR/IL-31R-mediated human iPSCSN activation.

**Figure 4:**
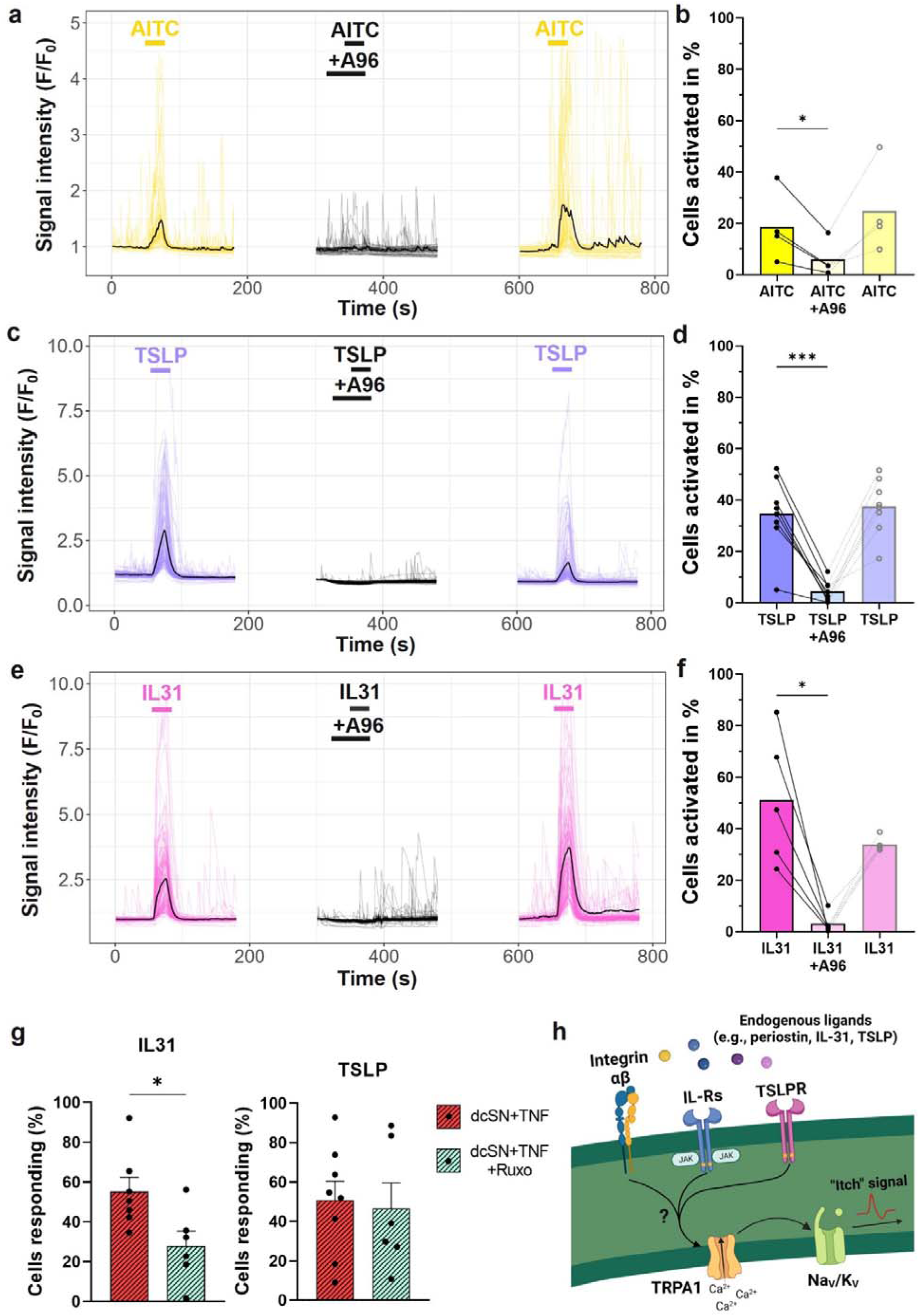
Contribution of TRPA1 and JAK1/2 to pruritogen sensitivity of iPSCSNs in the AD itch model. (a) Representative Ca^2+^-imaging of dcSN+TNF treated with 20s AITC (100 µM), 260s wash, 40s A967079 (10 µM) incl. 20s AITC+A967079, 280s wash and 20s AITC. (b) Percentage of cells reacting to AITC±A967079 (paired t test between initial AITC and +A96, n=4). (c) Representative Ca^2+^-imaging of dcSN+TNF treated with 20s TSLP (2 µg/mL), 260s wash, 40s A967079 (10 µM) incl. 20s TSLP+A967079, 280s wash and 20s TSLP. (d) Percentage of cells reacting to TSLP±A967079 (paired t test between initial TSLP and +A96, n=8). (e) Representative Ca^2+^-imaging of dcSN+TNF treated with 20s IL-31 (25 µg/mL), 260s wash, 40s A967079 (10 µM) incl. 20s IL-31+A967079, 280s wash, and 20s IL-31. (f) Percentage of cells reacting to IL-31±A967079 (paired t test between initial IL-31 and +A96, n=5). (g) Percentage of dcSN+TNF±Ruxolitinib (500 nM, 48h application) responding to IL-31 and TSLP (unpaired t test, n≥6). (h) Graphical summary of a proposed itch signaling mechanism via TRPA1. Data are presented as mean ± SEM, data points are biological replicates. (*=p<0.05; ***=p<0.001)

JAK1/2 inhibitors reduce IL-31R/TSLPR downstream signaling and have recently been shown to reduce pruritus in AD (Blauvelt et al., 2023). We therefore applied the JAK1/2 inhibitor ruxolitinib (500 nM) to dcSN+TNF cultures for 48h. Ruxolitinib-treated cells (dcSN+TNF+Ruxo) showed significantly lower responses to IL-31, but TSLP-sensitivity was unaffected (Figure 4g).

In summary, results suggest a potential pruritogen signaling pathway that revolves around the co-activation of TRPA1 for itch signal propagation in human iPSCSNs as has been shown in rodents (Figure 4h).

## DISCUSSION

Bidirectional interaction between skin cells and sensory neurons is an essential part of skin homeostasis and disease. For example, sensory neurons release neurotransmitters that induce keratinocyte proliferation (Roggenkamp et al., 2013), whereas keratinocytes produce nerve-sensitizing mediators (Talagas et al., 2020, Xu et al., 2022, Lin et al., 2024, Smith et al., 2025). To recreate this relationship, there have been attempts at sensory neuron-keratinocyte co-culture, using low Ca^2+^-containing medium to enable keratinocyte proliferation (Ulmann et al., 2007, Pereira et al., 2010). However, these studies did not physically separate both compartments and therefore keratinocytes grew alongside neuronal somata, rather than neurites. Campenot chambers (Campenot et al., 2009) have been utilized to create co-cultures similar to the model presented here, but have relied on animal-derived sensory neurons (Roggenkamp et al., 2012).

Previous iPSC-based co-culture approaches highlighted the potential of replacing animal-derived sensory neurons with patient-/disease-relevant cells (Guimaraes et al., 2018, Belamadni et al., 2022). However, these iPSC-based studies examined cells that either heavily clustered or were cultured in a closed environment, which prevented functional analysis, such as Ca^2+^-imaging and electrophysiology. Although iPSCSN incorporation into a skin model has been previously described (Muller et al., 2018), the complexity of compact 3D skin models means that neurons are inaccessible for functional testing. Meanwhile, the unique “deconstructed skin” setup described here made direct access to neurons possible, while allowing them to interact with the skin cells.

Comparing electrophysiological properties of 8-week iPSCSNs (Table 1) with published data (McDermott et al., 2019, Meents et al., 2019), McDermott et al. report a resting membrane potential of approximately-60 mV, we observe-52.2 mV, and Meents et al. see-39.3 mV. Furthermore, McDermott et al. showcase a general multi-AP firing ability, which we also observed (Fig. S2), whereas Meents et al. report only 3/22 iPSCSNs firing repeatedly. McDermott et al. describe a higher rheobase (∼100 pA) than our observation (∼12 pA), which is skewed by spontaneous AP-firing cells having a rheobase of 0 pA (excluding spontaneously firing cells, the rheobase was ∼39 pA). The preponderance of iPSCSN spontaneous firing we observed was also reported in 8-week Axol (Odawara et al., 2022) and other 7-week iPSCSNs (Hirano et al., 2021). In general, the iPSCSNs described in this study featured relevant electrophysiological properties of sensory neurons that have been described in other studies. However, between studies, there are many differences in iPSCSN sources and differentiation protocols that likely account for differences reported.

Even using the same differentiation protocol, a common issue with iPSCSN culture is inter-experimental and lot/donor variability (Schwartzentruber et al., 2018), which was also observed here with mATP responses varying from 0% to 78% iPSCSNs (Figure S1g). Furthermore, co-culture development introduces the variability of HEK donors (e.g. biopsy site, age, genetic background etc.). Despite this, we consistently observed a robust increase in TRPA1 expression and function in dcSN(+TNF) cultures and similar trends for increased sensitivity to the pruritogens TSLP and IL-31. Fittingly, TNF has also been shown to increase TRPA1 expression in non-neuronal cells (El Karim et al., 2015, Luostarinen et al., 2021). Moreover, responses to TSLP and IL-31 were inhibited by the TRPA1 inhibitor A967079, indicating that this ion channel mediates TSLPR and IL-31R Ca^2+^-signaling in human cells.

Until now, this has so far only been demonstrated in animal models (Wilson et al., 2013b, Cevikbas et al., 2014). Interestingly, more TSLP and IL-31 responding iPSCSN were blocked by TRPA1 inhibition than responded directly to AITC (Figure 4b/d/f). One potential explanation is that TRPA1-mediated responses to TSLP and IL-31 result from amplified microdomain signaling, allowing TRPA1-dependent signaling in response to TSLP and IL-31, but not AITC, in some cells.

By contrast, ruxolitinib pre-incubation only reduced sensitivity to IL-31. TSLP-derived signals appeared unaffected by this JAK inhibitor, in line with a reported phospholipase-C-dependent communication of TSLPR to TRPA1 (Wilson et al., 2013b). Potentially, ruxolitinib affected prior JAK-based signaling exchange in HEK-neuron communication during incubation time, rather than blocking acute Ca^2+^ signaling.

Overall, the human cell-based model described here enables investigation of itch signaling mechanisms and pharmacological testing of anti-pruritic substances. This valuable tool for AD research could also be adapted to other skin diseases e.g. by addition of other cytokines or incorporation of further skin/immune cells.

## MATERIAL & METHODS

See supplementary methods for more details, experimental setup and analysis.

### Tissue acquisition

All tissues used were acquired through commercial suppliers (Alphenyx, Marseille, France; AnaBios, San Diego, CA, USA) in charge of ethical agreements and informed donor consent in accordance with the declaration of Helsinki.

### Cell culture

Human epidermal keratinocytes (HEK) were isolated from surgical skin (Alphenyx, Marseille, France). Skin strips were digested with dispase II, epidermis separated, incubated in Trypsin/EDTA and HEK isolated through a cell strainer. For expansion, HEK were cultured in Epilife+HKGS (Gibco).

iPSCSN (Axol Bioscience, Cambridge, UK) were cultured in maintenance medium on 12 mm coverslips (Corning, NY, USA) for 8-weeks including mitomycin C (Merck) treatment on day 3. For contactless medium-based coculture (medSN), after 14-days HEK seeded in 24-well plate culture inserts were added on top of iPSCSN coverslips for further 21-day culture with 1:1 iPSCSN/HEK medium. For direct-contact coculture (dcSN), iPSCSN were cultured in the middle of 35 mm petri dishes for 14-days before seeding HEK in the perimeter and culturing for 14-days in 1:1 iPSCSN/HEK medium. Medium was then exchanged for 7-days culture in 1:1 iPSCSN/3DGro^TM^ differentiation medium.

Later models included cytokine treatment or addition of Th2 cell supernatant for 48-hours before experimental readout.

### Immunofluorescence

Cell cultures or frozen biopsy sections were washed with PBS, fixed with 4 % paraformaldehyde for 20 min and washed again. After 1 h of blocking/permeabilization (0.2 % Triton X-100 in Seablock/Superblock, ThermoFisher), primary antibodies were added (see Table S2) and incubated overnight at 4 °C. Consecutive washing was followed by secondary antibody incubation for 3-4 h at 4 °C. Slides were washed in PBS and mounted with ProLong Glass (ThermoFisher).

### Gene expression experiments

Cells were lysed in Cells-to-CT lysis solution for 1-step gene expression experiments (C2CT, Invitrogen/ThermoFisher) according to manufacturer’s instructions. Lysate was mixed with Taqman 1-step qRT-PCR Mix and applied to custom-made gene expression cards (Applied Biosystems/ThermoFisher). Gene expression was quantified in duplicate, compared to 2 housekeeping genes (glyceraldehyde-3-phosphate dehydrogenase, GAPDH and hypoxanthine phosphoribosyltransferase 1, HPRT1) and normalized to iPSCSNs at d0.

### Ca^2+^-imaging

Intracellular [Ca^2+^] was visualized by 488 nm excitation of Calbryte^TM^ 520 AM (5 µg/mL, AAT Bioquest, Pleasanton, CA, USA) and 1x PowerLoadTM (ThermoFisher) for 30 min and consecutive 15 min washout/equilibration with extracellular solution (ECS, Brainphys imaging optimized medium, STEMCELL). Fluorescence intensity was measured over 180 s at 1 frame/s. Compounds were applied for 20 s with 300 s wash period in between: capsaicin (1 µM, Sigma-Aldrich), menthol (250 µM, Alfa Aesar), AITC (100 µM, Merck), mATP (100 µM, Sigma-Aldrich), TSLP (2 µg/mL, Miltenyi), IL-31 (25 µg/mL, STEMCELL), periostin (25 µg/mL; Biolegend), KCl (50 mM, Sigma-Aldrich). Where applicable, ruxolitinib (500 nM, InvivoGen) was added to cells with cytokines for 48h, A967079 (10 µM, Tocris Bioscience) was applied during compound perfusion experiments.

### Patch-clamp electrophysiology

Recordings were made as described before (Chakrabarti et al., 2018). For analysis, cells had to show a resting membrane potential (Rm) <-40 mV and respond to current injections with an evoked action potential (AP).

## DATA AVAILABILITY

No large datasets were generated or analyzed in this study. Experimental data can be requested from the corresponding author. (Placeholder: Raw data will be available from the University of Cambridge Apollo repository following publication.)

## ACKNOWLEDGEMENTS

We want to thank Nicholas Holzscheck for his help with creating a Ca^2+^-imaging analysis script, as well as Simona Lange and Anna Geueke for helpful project discussions and input regarding experiments. Additionally, we thank Julia Korn and Heiko Mielke for help with cell culture and Luke Pattison and Alex Cloake for guidance on electrophysiology.

Figures 1a, 2a/c, 3d and 4h (the graphical abstract) were created using Biorender.com.

## AUTHOR CONTRIBUTIONS

HM: Conceptualization, Data Curation, Formal Analysis, Investigation, Methodology, Visualization, Writing – Original Draft Preparation

BA: Methodology, Writing – Review and Editing

HR: Conceptualization, Funding Acquisition, Writing – Review and Editing

JS: Conceptualization, Project Administration, Resources, Supervision, Writing – Review and Editing

EStJS: Conceptualization, Project Administration, Resources, Supervision, Writing – Review and Editing

## Abbreviations

AD: atopic dermatitis;
AP: action potential
dcSN: direct contact sensory neuron co-culture with HEK
HEK: human epidermal keratinocytes
iPSCSN: human induced pluripotent stem cell-derived sensory neurons;
TNF: Tumor necrosis factor (alpha)
TSLP: Thymic stromal lymphopoietin;
TRPV1/TRPM8/TRPA1: Transient receptor potential cation channel subfamily V1/M8/A1

## SUPPLEMENTARY INFORMATION

**Supplementary Figure S1:**
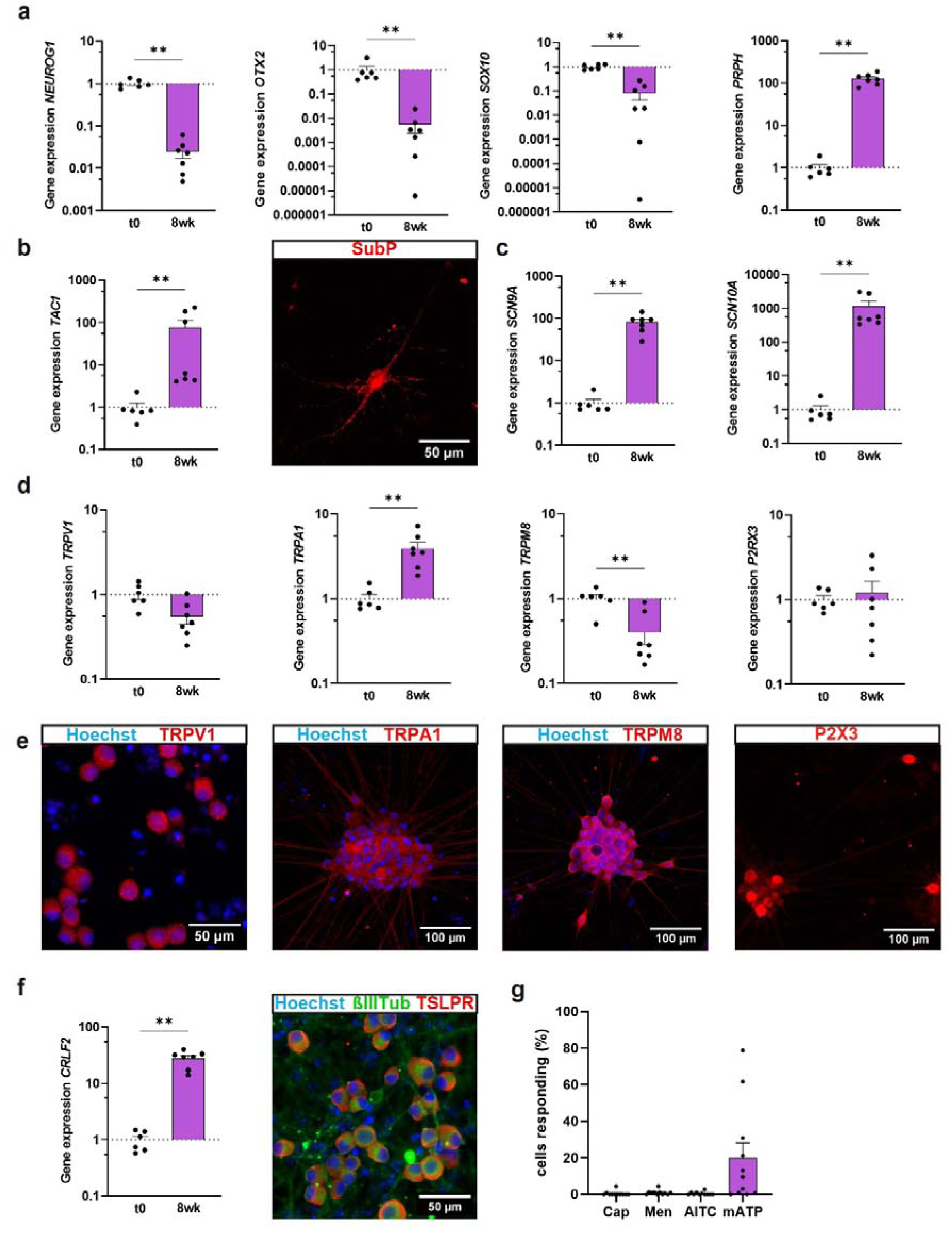
Gene and protein expression data for iPSCSN characterization. (a) Expression of progenitor-and stem cell-characteristic genes *NEUROG1*, *OTX2*, and *SOX10*, as well as neurofilament peripherin (*PRPH*) in mature iPSCSNs (8wk) compared and normalized to progenitors (t0). (b) Gene expression (*TAC1*) and immunocytochemistry (SubP, red) of neurotransmitter Substance P. (c) Gene expression of voltage-gated sodium channel subunits Nav1.7 (*SCN9A*) and Nav1.8 (*SCN10A*). (d) Gene expression and (e) immunocytochemistry (red; nuclei/Hoechst, blue) of cation channels TRPV1 (*TRPV1*), TRPA1 (*TRPA1*), TRPM8 (*TRPM8*) and ATP-purinoceptor P2X3 (*P2RX3*). (f) Gene expression (*CRLF2*) and immunocytochemistry (red) of itch and atopic dermatitis-associated receptor TSLPR in combination with beta-III-Tubulin (βIIITub, green) and nuclei (Hoechst, blue). (g) Bar graph showing the percentage of mature iPSCSNs reacting to capsaicin (Cap, 1 µM), menthol (Men, 250 µM), allyl isothiocyanate (AITC, 100 µM), α,β-methylene ATP (mATP, 100 µM) in Ca^2+^-imaging experiments (KCl-positive-only (50 mM), n=11, 1834 cells). Gene expression data are presented as mean ± SEM, relative fold-change to t0, 2^-ΔΔCt^, biological replicates n≥4 for t0, n≥6 for 8wk; Mann-Whitney-test, **=p<0.01. Scale bars represent 50/100 µm (indicated).

**Supplementary Figure S2:**
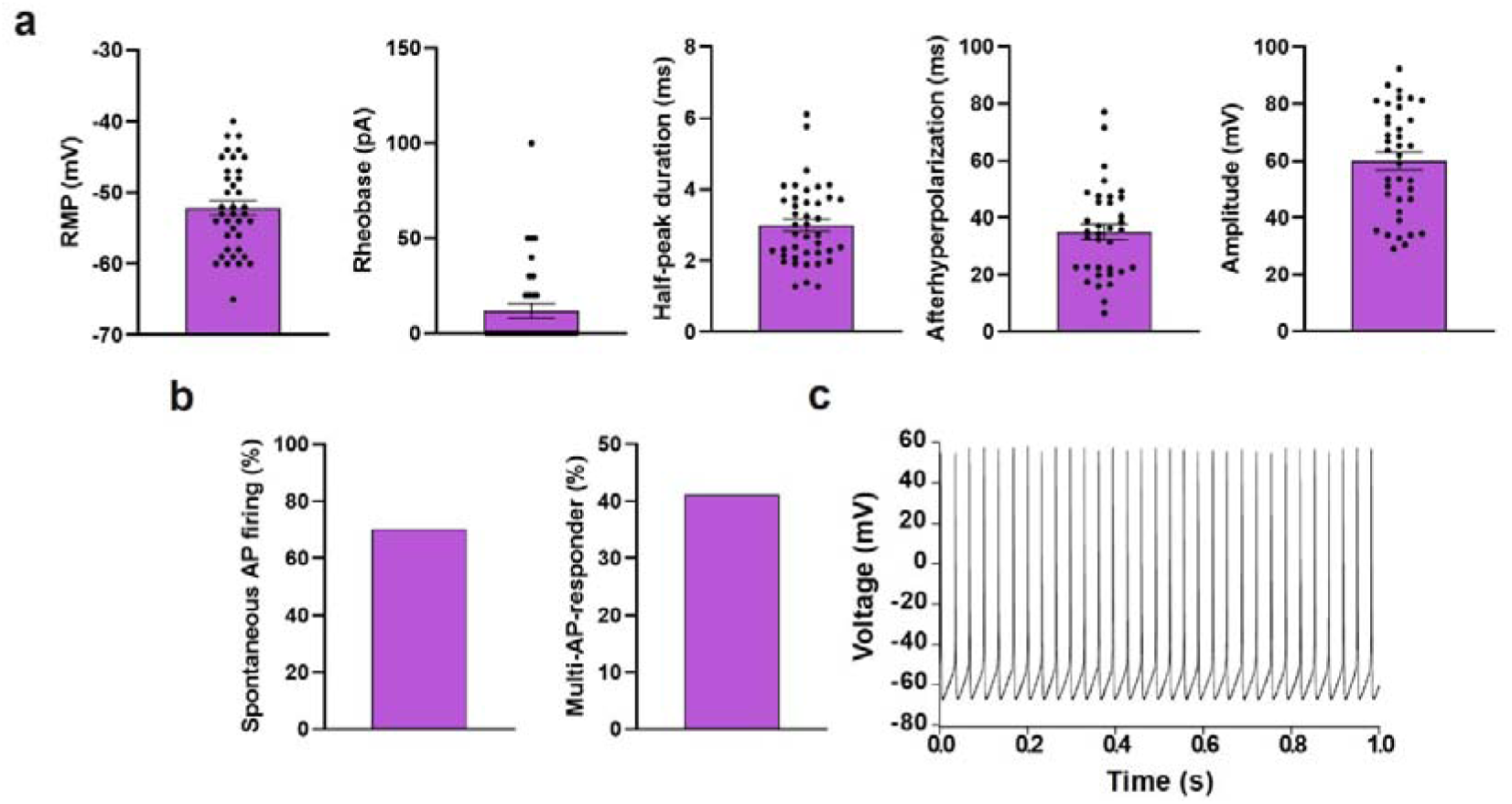
Electrophysiology parameters of iPSCSNs. (a) Resting membrane potential (RMP) of analyzed cells and action potential parameters AP threshold/rheobase (spontaneous firing cells = 0 pA), half-peak duration, afterhyperpolarization time and peak amplitude. (b) Percentage of spontaneous AP-firing cells without current injection and multi-AP responders in reaction to ascending currents above rheobase. (c) Patch-clamp trace after cell access (without current injection) illustrates the ability of iPSCSNs to fire spontaneous action potentials. Data are presented as mean ± SEM (n≥36, individual cells from 7 separate cultures).

**Supplementary Figure S3:**
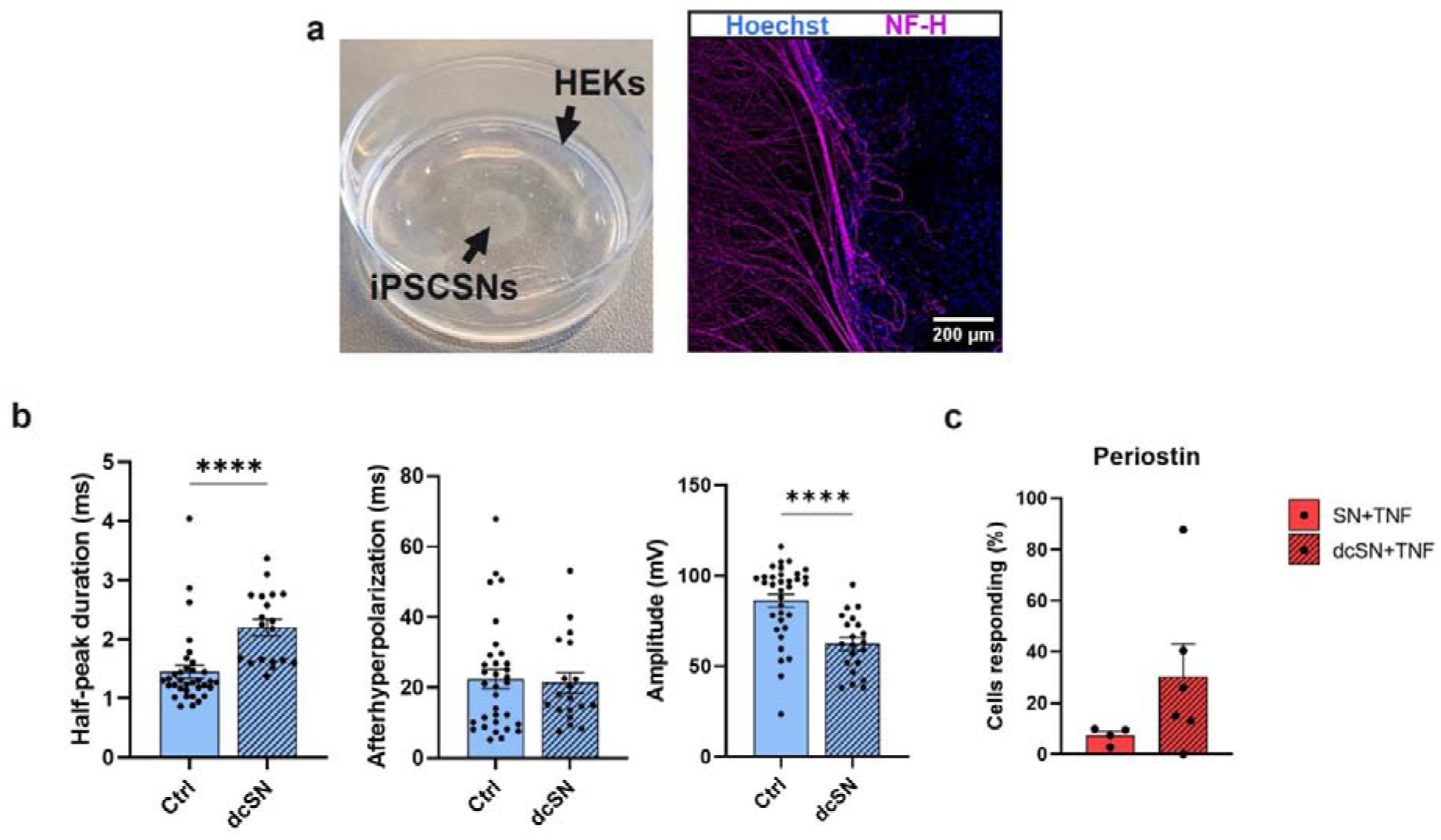
Additional data for iPSCSN co-culture. (a) Differentiated dcSN culture in the middle of a petri dish with bulging HEKs in the outer perimeter and immunohistochemistry staining visualizing the connection between neurites (NF-H, purple) and keratinocytes (nuclei/ Hoechst, blue) (Scale bar represents 200 µm). (b) Comparison of dcCtrl and dcSN action potential parameters half-peak duration, afterhyperpolarization time and peak amplitude (Mann-Whitney-test, n≥32/20 (Ctrl/dcSN) cells from 5/4 individual cultures). (c) Comparison of TNF-treated SN or dcSN responding to Periostin (KCl-positive-only, unpaired t test, data points are biological replicates). Data are presented as mean ± SEM. (****=p<0.0001)

## SUPPLEMENTARY METHODS

### Tissue acquisition

All tissues used were acquired through commercial suppliers in charge of ethical agreements and informed donor consent in accordance with the declaration of Helsinki.

Primary human skin was obtained from healthy consented donors undergoing plastic surgery (Alphenyx, Marseille, France) in accordance with the declaration of Helsinki. For skin cell dissociation and tissue integrity, procurement, serology testing and shipping, the timeline was strictly set to 24h from the time of tissue removal.

**Supplementary Table S2:** Donor information for skin used for keratinocyte extraction, as well as immunohistochemistry.

| Sex | Age | Area |
| --- | --- | --- |
| M | 51 | Face |
| F | 23 | Breast |
| F | 31 | Breast |
| F | 28 | Abdomen |
| F | 62 | Breast |
| M | 29 | Foreskin |

Human dorsal root ganglia (DRG) were acquired from AnaBios (San Diego, CA, USA), sourced from fully consented organ donors including medical review.

### Skin cell dissociation and culture

Primary skin cells were isolated from healthy surgical skin tissue. Skin was cut into 3-5 mm wide strips and digested using dispase II solution (2.4 U/mL in HBSS; Sigma-Aldrich, St. Louis, MO, USA) for 3 hours at 37 °C or overnight at 4 °C. Epidermis was mechanically separated from dermis using forceps and incubated in trypsin/EDTA (37 °C, 5 min; Bioconcept, Allschwil Switzerland). Human epidermal keratinocytes (HEK) were isolated by grinding resulting epidermal pieces through a cell strainer (70 µm) and rinsing with stopping solution, which consists of 10% fetal calf serum (heat-inactivated FCS; Sigma-Aldrich) + 1% penicillin/streptomycin (Gibco/Thermo Fisher Scientific, Waltham, MA, USA) in phosphate buffered saline (PBS, Gibco). After centrifugation (500×g, 5 min) cells were resuspended in culture medium (EpiLife + HKGS, Gibco; 1% penicillin/streptomycin). Dermis was minced and incubated with collagenase type I (312 U/mL, Gibco) for 2 hours (37 °C) to enable human dermal fibroblast (HDF) isolation, which was conducted as for keratinocytes, but with DMEM medium (+10 % fetal calf serum, 1% penicillin/streptomycin, 1% GlutaMAX, Gibco). Cells were plated and grown to ∼80 % confluency, at which point they were washed with PBS and incubated in trypsin/EDTA (37 °C, 5 min) for detachment, trypsin was inactivated using the respective cell type medium including 10 % FCS. Cells were either passaged for extended culture or taken up in freezing medium (culture medium +20% DMSO, Sigma-Aldrich) for storage in liquid nitrogen. Cells were counted using EVE (NanoEnTek, Seoul, Korea) and 10^6^ cells were either stored or seeded into a new T175 flask. Maximum passage number for experimental use of both keratinocytes and fibroblasts was P4.

### Induced pluripotent stem cell-derived sensory neurons (iPSCSNs)

IPSCSNs (Axol Bioscience, Cambridge, UK; ax0055, male progenitor sensory neurons) were of CD34+-cord blood cell origin and reprogrammed using a non-integrating episomal vector system.

12 mm coverslips (Corning® BioCoat® Poly-D-Lysine-treated, Corning, NY, USA) or 35 mm petri dishes (Greiner Bio-one, Kremsmünster, Austria) were freshly coated with Poly-D-lysine/H_2_O solution (PDL, 0.1 mg/mL, 500 µL per well; Merck, Darmstadt, Germany), incubated for 1 hour at room temperature (RT, 21 °C), washed twice with ddH2O and dried for 1-2 hours at RT. Next, coverslips were coated with Surebond XF solution (Axol, 1:200 in PBS) for 3 hours at 37 °C, followed by Growth factor-reduced Matrigel (Corning) 1:100 in PBS for 60 min at 37 °C followed. Without washing, iPSCSNs were seeded at 200,000 cells/mL in Axol plating medium (ax0035) and incubated at 37 °C/ 5% CO2 overnight. Cells were maintained in maintenance medium (Axol, ax0060, mixed 1:1 with Schwartzentruber P2 (Schwartzentruber et al., 2018) based on Neurobasal Plus and B27 Plus (Gibco); 25 ng/mL NGF/BDNF/GDNF/NT-3, 200 µM or 35 µg/mL ascorbic acid). Maturation maximizer (Axol, ax0058) was freshly added 1:100 to culture medium for 8 weeks with half medium changes every 3 days. During the first week, DAPT (N-[N-(3,5-difluorophenacetyl)-L-alanyl]-S-phenylglycine t-butyl ester, 1 µM, Sigma-Aldrich) was added to the culture medium to induce differentiation and limit extensive cell clustering (Saito-Diaz et al., 2021). To eliminate non-differentiating, proliferating cells from the culture, cells were treated with filter-sterilized mitomycin C (2 µg/mL, in maintenance medium; Sigma-Aldrich) for 2 h at 37 °C on day 3 after plating. After treatment, cells were washed once with maintenance medium and culturing continued. Laminin (500 ng/mL, Merck) was added to the maintenance medium for 2 weeks ahead of planned experiments to avoid detachment/peeling. For a visualisation of the differentiation protocol, see Figure 1a.

### Co-Culture

For contactless co-culture with shared medium (medSN), iPSCSNs (200,000 cells) were plated on coverslips and cultured for 2 weeks. Thereafter, 24 well plate cell culture inserts were coated with 100 µL human collagen I (50 µg/mL, Santa Cruz, Dallas, TX, USA) for 2 hours at 37 °C. Without washing, HEK were seeded into the inserts (50,000 cells), cultured for 1 day in EpiLife+HKGS and transferred to the iPSCSN wells. MedSN cells were cultured with a mixture of 1:1 HEK/iPSCSN maintenance medium for 6 days in submerged culture and for 2 weeks at air-liquid interface (350 µL in-well medium volume) with medium changes every 2 days. SN medCtrl were treated the same way except for adding HEK. For a visualisation of the setup, see Figure 2a.

For co-culture with direct contact between cell types (dcSN), iPSCSNs (100,000 cells) were plated in the middle of 35 mm petri dishes (pre-coated as described for coverslips in the previous section) using 10×10 mm cloning cylinders (C2059, Sigma-Aldrich). Cylinders were removed after 18 hours and cells were cultured for 2 weeks as described above. Afterwards,

HEK (100.000 cells) were seeded into the outer perimeter of iPSCSN-containing petri dishes using customised glass inserts (22 mm outer, 18.4 mm inner diameter, 12 mm height; Hilgenberg, Malsfeld, Germany). Glass inserts were removed after 18 hours and cells were cultured with a mixture of 1:1 HEK/iPSCSN maintenance medium for 2 weeks, changing medium every 2 days. HEK maintenance medium was then exchanged for 3DGro^TM^ Skin Differentiation Medium (Merck) for 1 week. For a visualisation of the setup, see Figure 2c and S3a.

Cytokine treatment with IL-4/IL-13 (50 ng/mL; STEMCELL Technologies, Vancouver, Canada) or TNF-α (20 ng/mL; STEMCELL) was done for 48 hours, reapplication followed after 24h. For inhibitor tests, ruxolitinib (500 nM, InvivoGen, San Diego, CA, USA) was added to cells together with cytokines for 48h, A967079 (10 µM, Tocris Bioscience, Bristol, UK) was applied alongside tested substances (see Ca^2+^-imaging section).

### Th2 cells

Th2 cell supernatant: Th2 cells were induced by isolating naïve CD4+ T cells from peripheral blood mononuclear cells (PBMCs, EasySepTM Human Naïve T cell isolation kit, STEMCELL) and polarizing for one week using anti-CD2/CD3/CD28 MACSiBeads (Miltenyi Biotec Bergisch Gladbach, Germany), IL-4 (50 ng/ml, STEMCELL) and anti-IFNγ antibodies (5 µg/ml, BioLegend, San Diego, CA, USA).

Th2 supernatant was collected, pooled, added to a 1 kDa molecular weight cut-off column (Pall Corporation, NY, USA) and centrifuged (3h, 3000×g). Retained volume (∼150 µL) was added to culture, where indicated, for 48 hours (re-application after 24 hours) before experiments were conducted.

### Immunohistochemistry / Immunocytochemistry

Biopsies were embedded in optimal cutting temperature compound (Sakura Tissue-Tek O.C.T.; Alphen aan den Rijn, Netherlands), frozen, cut on a cryostat (Leica, Wetzlar, Germany) in 14 µm sections, placed on microscope slides and dried. Using a Coplin jar, slides were first soaked in PBS (without Ca^2+^/Mg^2+^), then fixed with cold paraformaldehyde (4 %) for 20 min and washed three times with PBS. Blocking buffer (1:1 combination of Sea Block, ThermoFisher; Superblock, ThermoFisher; + 0.2 % Triton X-100) was directly applied to samples for 1 hour and then exchanged for primary antibody solution (e.g. NF-H 1:500 in blocking buffer), and incubated overnight at 4 °C. Slides were rinsed three times with PBS and a solution of fluorochrome-labelled secondary antibody (1:800, incl. Hoechst 1:2000) was applied for 3-4 hours at 4 °C. Following 3 washes with PBS, slides were mounted with ProLong Glass and left to cure.

Immunocytochemistry with iPSCSNs followed the same protocol as IHC, but with the process conducted in dishes and cells topped with a coverslip, or cells were grown on coverslips and treated in well plates before mounting on glass slides.

**Supplementary Table S3:** Primary antibody list.

| **Antigen** | **Species** | **Conc.** | **Supplier** | **Cat. No.** |
| --- | --- | --- | --- | --- |
| **NF-H** | Chicken | 1:2000 | Abcam | ab4680 |
| **P2X3** | Rabbit | 1:200 | Alomone | APR-016 |
| **Peripherin** | Rabbit | 1:200 | Abcam | ab4666 |
| **Substance P** | Guinea | 1:50 | Novus bio | NB300-187 |
| **TRPA1** | Rabbit | 1:200 | Invitrogen | PA5-34282 |
| **TRPM8** | Sheep | 1:200 | Invitrogen | OST00028W |
| **TRPV1** | Rabbit | 1:200 | Abcam | ab3487 |
| **TSLPR** | Goat | 1:200 | R&D | AF981 |
| **β-III-Tubulin** | Rabbit | 1:400 | Abcam | ab18207 |

**Supplementary Table S4:** Secondary antibody list.

| **Host** | **Anti-species** | **Alexa Fluor** | **Supplier** | **Cat. No.** |
| --- | --- | --- | --- | --- |
| **Donkey** | Chicken | 488 | Dianova | 703-546-155 |
| **Donkey** | Goat | 647 | Dianova | 705-175-003 |
| **Donkey** | Guinea pig | 647 | Abcam | ab150187 |
| **Donkey** | Rabbit | 488 | Invitrogen | A21206 |
| **Donkey** | Rabbit | 546 | Invitrogen | A10040 |
| **Donkey** | Sheep | 546 | Invitrogen | A21098 |

### Ca^2+^-imaging

The perfusion setup for Ca^2+^-imaging was based on a VCS-8-Pinch computer-controlled Valve Control System combined with a micromanipulator, multi-barrel perfusion pencil (AutoMate Scientific, Berkeley, CA, USA), QE-1 Quick Exchange Platform (Multi Channel Systems, Reutlingen, Germany) and a Preciflow pump (Lambda Instruments, Baar, Switzerland). The Ca^2+^ indicator Calbryte^TM^ 520 AM (5 µg/mL in-well concentration, AAT Bioquest, Pleasanton, CA, USA) was used together with 1x PowerLoad^TM^ (ThermoFisher), enabling the use of maintenance medium for cell loading, substance dilution and washing steps. BrainPhys imaging optimized medium (STEMCELL) was used as extracellular solution (ECS) and for compound uptake/dilution if not stated otherwise.

Intracellular Ca^2+^ concentration changes were visualized by 488 nm excitation following the uptake of Calbryte 520 AM (30 min incubation, 37 °C, wash out 2x, replaced with ECS, 15 min resting time). Fluorescence intensity was measured over 180 s at a frequency of 1 frame/s. Depending on the experiment, a selection of the following compounds were applied for 20 s with 300 s wash period in between: capsaicin (1 µM, 4 mM stock in 100% ethanol; Sigma-Aldrich), menthol (250 µM, 1 M stock in 100% ethanol; Alfa Aesar, Ward Hill, MA, USA), AITC (100 µM, 400 mM stock in 100% ethanol; Merck), mATP (100 µM, Sigma-Aldrich), TSLP (2 µg/mL, Miltenyi), IL-31 (25 µg/mL, STEMCELL), periostin (25 µg/mL; Biolegend, San Diego, CA, USA), KCl (50 mM, Sigma-Aldrich). For inhibitor experiments with A967079, the previous substances were applied simultaneously with the inhibitor from a single tube to avoid mixing effects.

Timeseries recordings were loaded into ImageJ/Fiji, regions of interest (ROI) were marked based on positive control responses and, if necessary, adjusted using a brightfield image. The mean grey value (fluorescence intensity) was then measured for each individual recording and ROI. For analysis, a custom script was written in R. In short, measurements were background corrected (to a ROI without cells) and each cell’s fluorescence intensity normalized to its baseline value (average of the first 9 seconds of recording prior to any stimulation); cells were omitted from further analysis if negative signals <-5 were detected. A rolling mean (5 s) threshold for activation by respective compounds was based on 50x baseline standard deviation to avoid spontaneously active cells from being detected as false positive responders. For responder statistics, only cells responding to KCl were included.

### Patch-clamp electrophysiology

Patch-clamp recordings were made for a maximum of 3-hours from one dish. BrainPhys imaging optimized medium was used as extracellular solution (ECS). Patch pipettes of 4–9 MΩ were pulled with a P-97 Flaming/Brown puller (Sutter Instruments; Novato, CA, USA) from borosilicate glass capillaries and the intracellular solution used contained (in mM): KCl (110), NaCl (10), MgCl_2_ (1), EGTA (1), HEPES (10), Na_2_ATP (2), Na_2_GTP (0.5) adjusted to pH 7.3 with KOH. Osmolarity was approximately 320±10 mOsm. Recordings were made using an EPC-10 amplifier (HEKA, Lambrecht, Germany) together with HEKA Patchmaster software. For voltage clamp experiments, iPSCSNs were held at −60 mV. Only neurons where an action potential (AP) could be evoked in response to current injections and that showed a resting membrane potential more negative than −40 mV were analysed. Images of neurons were captured using a 40× objective on a Nikon Eclipse Ti-S microscope (Tokyo, Japan) and a Zyla 5.5 sCMOS camera (Andor, United Kingdom).

Electrophysiological recordings were analysed using HEKA Fitmaster Software and Wavemetrics Igor Pro (Portland, OR, USA). Characteristic features of recorded APs were resting membrane potential (Rm), rheobase (AP threshold), half peak duration (HPD), afterhyperpolarization time (AHP), amplitude; spontaneous firing and/or the response of multiple APs to stimulation (Figure 2g and Figure S2a/b). Additionally, whether or not an inflection/“hump” occurred during the repolarisation phase was analysed by examining each AP’s derivation plot, such a hump being considered characteristic of nociceptors (Djouhri et al., 1998, Davidson et al., 2014).

### Cell culture tracking and fluorescence imaging

Cell culture images were taken with an EVOS XL Imaging system (Life technologies/ThermoFisher). Fluorescent images and Ca^2+^-recordings were obtained using either a Zeiss Axio Observer Z1 epifluorescence microscope and AxioCam 503 mono camera with Zen 3.0 software (Oberkochen, Germany) or Leica TCS SP5 II confocal microscope (using HyD detectors) with LAS X Software. Z-Stacks of tissues or cells were imaged with the Leica TCS SP5 II confocal microscope and processed to result in a Maximum Intensity Projection. Confocal images were taken in a sequential channel-separated scanning mode without simultaneous excitation of different wavelengths and an automatic wavelength-specific dye separation was performed to avoid cross-reactions between fluorophores.

### RT-qPCR Gene expression cards

1-step C2CT mix was freshly prepared (100 µL per iPSCSN sample) from C2CT lysis solution (Invitrogen), 1:100 RNAse inhibitor (Applied Biosystems/ThermoFisher), 1:100 DNase I (Invitrogen). Cells were washed with PBS, incubated with C2CT mix for 5-7 minutes and pipetted up and down until lysed completely. In case of dcSN co-cultures, iPSCSN cell bodies first needed to be excised with a small pipette tip for lysis. Afterwards, 1:10 C2CT Stop solution (Invitrogen) was applied, mixed well, incubated for 2-3 minutes and stored at - 80 °C. 11 µL lysate was mixed together with 27.5 µL Taqman 1-step qRT-PCR Mix and 71.5 µL nuclease-free water, and applied to custom-made TaqMan gene expression cards (Applied Biosystems). Experiments were performed using a 7900HT Fast Real-Time PCR System (Applied Biosystems) according to the manufacturer’s instructions. Gene expression per sample was quantified in duplicate, compared to 2 housekeeping genes (glyceraldehyde-3-phosphate dehydrogenase, GAPDH and hypoxanthine phosphoribosyltransferase 1, HPRT1) based on their suitability for PCR experiments with neuronal tissue (Jiang et al., 2020) and normalized to the mean of all iPSCSN lots at the day of plating (d0, progenitor cells). Resulting ΔΔC_T_ values were displayed as relative target number (2^-ΔΔC^, relative gene expression to d0 with mean=1).

**Supplementary Table S5:** TaqMan probes used for gene expression cards in RT-qPCR experiments.

| Assay ID | Gene symbol(s) | Gene Name(s) | Species | Amplicon Length |
| --- | --- | --- | --- | --- |
| Hs01029249_s1 | *NEUROG1* | neurogenin 1 | Human | 67 |
| Hs00222238_m1 | *OTX2* | orthodenticle homeobox 2 | Human | 72 |
| Hs00366918_m1 | *SOX10* | SRY-box 10 | Human | 102 |
| Hs01021970_m1 | *RUNX1* | runt related transcription factor 1 | Human | 77 |
| Hs00196608_m1 | *PRPH* | peripherin | Human | 83 |
| Hs01021011_m1 | *NTRK1* | neurotrophic receptor tyrosine kinase 1 | Human | 62 |
| Hs01076699_m1 | *SCN9A* | sodium voltage-gated channel alpha subunit 9 | Human | 69 |
| Hs01045137_m1 | *SCN10A* | sodium voltage-gated channel alpha subunit 10 | Human | 70 |
| Hs01125554_m1 | *P2RX3* | purinergic receptor P2X 3 | Human | 84 |
| Hs01100741_m1 | *CALCA* | calcitonin related polypeptide alpha | Human | 84 |
| Hs99999905_m1 | *GAPDH* | glyceraldehyde 3-phosphate dehydrogenase | Human | 122 |
| Hs02800695_m1 | *HPRT1* | hypoxanthine phosphoribosyltransferase 1 | Human | 82 |
| Hs00243225_m1 | *TAC1* | tachykinin precursor 1 | Human | 78 |
| Hs00175798_m1 | *TRPA1* | transient receptor potential cation channel subfamily A member 1 | Human | 124 |
| Hs00218912_m1 | *TRPV1* | transient receptor potential cation channel subfamily V member 1 | Human | 94 |
| Hs01066596_m1 | *TRPM8* | transient receptor potential cation channel subfamily M member 8 | Human | 108 |
| Hs00965056_m1 | *IL4R* | interleukin 4 receptor | Human | 84 |
| Hs00609812_m1 | *IL13RA1* | interleukin 13 receptor subunit alpha 1 | Human | 106 |
| Hs00371172_m1 | *IL31RA* | interleukin 31 receptor A | Human | 78 |
| Hs00845692_m1 | *CRLF2* | cytokine receptor-like factor 2 | Human | 75 |
| Hs00608346_m1 | *F2RL1* | F2R like trypsin receptor 1 | Human | 119 |
| Hs00173590_m1 | *NPPB* | natriuretic peptide B | Human | 82 |
| Hs01042313_m1 | *TNFRSF1A* | TNF receptor superfamily member 1A | Human | 150 |
| Hs00961748_m1 | *TNFRSF1B* | TNF receptor superfamily member 1B | Human | 74 |
| Hs01099348_m1 | *TRPV4* | transient receptor potential cation channel subfamily V member 4 | Human | 65 |

## Statistical analysis

Data were analysed using GraphPad Prism 9.3.1 (San Diego, USA). Structurally different data (paired/unpaired, group size) were analysed using appropriate statistical test methods, indicated in the corresponding figure legends. Briefly, for comparisons of two groups, unpaired two-tailed t-tests were performed, Mann-Whitney test / ranked comparison for nonparametric data. When comparing 3 or more groups, ANOVA (one-way for single parameter, two-way for multiple parameter analysis, such as Ca^2+^-imaging responses) was followed by Tukey’s (One/Two-way ANOVA, all samples compared to each other), Dunnett’s (One/Two-way ANOVA, all samples compared to a control), Dunn’s (Kruskal-Wallis ranked comparison, all samples compared to each other or to a control) multiple comparisons test. Paired data was analysed using paired two-tailed t-test. Differences were considered significant if p < 0.05 (*=p<0.05; **=p<0.01, ***=p<0.001, ****=p<0.0001). For every experiment/graph, single data points were shown and therefore data distribution and dispersion were displayed. In addition, standard error of the mean was presented for an estimate of the sample mean accuracy.

## References

Al, B., Traidl, S., Holzscheck, N., Freimooser, S., MIEßner, H., Reuter, H., Dittrich□Breiholz, O., Werfel, T. & Seidel, J. A. 2024. Single□cell RNA sequencing reveals 2D cytokine assay can model atopic dermatitis more accurately than immune□competent 3D setup. Experimental Dermatology, 33.

Anand, U., Otto, W. R., Casula, M. A., Day, N. C., Davis, J. B., Bountra, C., Birch, R. & Anand, P. 2006. The effect of neurotrophic factors on morphology, TRPV1 expression and capsaicin responses of cultured human DRG sensory neurons. Neuroscience Letters, 399, 51–56.

Belamadni, A., Ren, D., Jayaraj, N., George, D., Miller, R. & Mencihella, D. 2022. Integrating Sensory Neurons with Keratinocytes to Model Painful Diabetic Neuropathy on a Chip. The Journal of Pain, 23, 10.

Berna, R., Mitra, N., Lou, C., Wan, J., Hoffstad, O., Wubbenhorst, B., Nathanson, K. L. & Margolis, D. J. 2021. TSLP and IL-7R Variants Are Associated with Persistent Atopic Dermatitis. J Invest Dermatol, 141, 446–450 e2.

Blauvelt, A., Kircik, L., Papp, K. A., Simpson, E. L., Silverberg, I., Jonathan, Kim, B. S., Kwatra, S. G., Kuligowski, M. E., Venturanza, M. E., Wei, S. & Szepietowski, J. C. 2023. Rapid pruritus reduction with ruxolitinib cream treatment in patients with atopic dermatitis. Journal of the European Academy of Dermatology and Venereology, 37, 137–146.

Campenot, R. B., Lund, K. & Mok, S.-A. 2009. Production of compartmented cultures of rat sympathetic neurons. Nature Protocols, 4, 1869–1887.

Cevikbas, F., Wang, X., Akiyama, T., Kempkes, C., Savinko, T., Antal, A., Kukova, G., Buhl, T., Ikoma, A., Buddenkotte, J., Soumelis, V., Feld, M., Alenius, H., Dillon, S. R., Carstens, E., Homey, B., Basbaum, A. & Steinhoff, M. 2014. A sensory neuron-expressed IL-31 receptor mediates T helper cell-dependent itch: Involvement of TRPV1 and TRPA1. J Allergy Clin Immunol, 133, 448–60.

Chakrabarti, S., Pattison, L. A., Singhal, K., Hockley, J. R. F., Callejo, G. & Smith, E. S. J. 2018. Acute inflammation sensitizes knee-innervating sensory neurons and decreases mouse digging behavior in a TRPV1-dependent manner. Neuropharmacology, 143, 49–62.

Charest, J. L., Jennings, J. M., King, W. P., Kowalczyk, A. P. & García, A. J. 2009. Cadherin-Mediated Cell–Cell Contact Regulates Keratinocyte Differentiation. Journal of Investigative Dermatology, 129, 564–572.

Danso, M. O., VAN Drongelen, V., Mulder, A., VAN Esch, J., Scott, H., VAN Smeden, J., EL Ghalbzouri, A. & Bouwstra, J. A. 2014. TNF-α and Th2 Cytokines Induce Atopic Dermatitis–Like Features on Epidermal Differentiation Proteins and Stratum Corneum Lipids in Human Skin Equivalents. Journal of Investigative Dermatology, 134, 1941–1950.

Donglang, G., Tongtong, L., Dan, C., Chan, Z., Changming, W., Guang, Y., Yan, Y. & Zongxiang, T. 2021. Comparative Study on Different Skin Pruritus Mouse Models. Front Med (Lausanne), 8, 630237.

EL Karim, I., Mccrudden, M. T., Linden, G. J., Abdullah, H., Curtis, T. M., Mcgahon, M., About, I., Irwin, C. & Lundy, F. T. 2015. TNF-alpha-induced p38MAPK activation regulates TRPA1 and TRPV4 activity in odontoblast-like cells. Am J Pathol, 185, 2994–3002.

Guimaraes, M. Z. P., De Vecchi, R., Vitoria, G., Sochacki, J. K., Paulsen, B. S., Lima, I., Rodrigues Da Silva, F., Da Costa, R. F. M., Castro, N. G., Breton, L. & Rehen, S. K. 2018. Generation of iPSC-Derived Human Peripheral Sensory Neurons Releasing Substance P Elicited by TRPV1 Agonists. Front Mol Neurosci, 11, 277.

Hirano, M., Huang, Y., Vela Jarquin, D., De La Garza Hernandez, R. L., Jodat, Y. A., Luna Ceron, E., Garcia-Rivera, L. E. & Shin, S. R. 2021. 3D bioprinted human iPSC-derived somatosensory constructs with functional and highly purified sensory neuron networks. Biofabrication, 13.

Jha, M. K., Han, Y., Liu, Z., Hara, Y., Langohr, I. M., Morel, C., Maloney, C. L., Piepenhagen, P., Xing, H., Bodea, C. A., Bangari, D. S., Mattoo, H. & Hicks, A. 2025. Type 2 cytokines pleiotropically modulate sensory nerve architecture and neuroimmune interactions to mediate itch. J Allergy Clin Immunol, 156, 1066–1081 e12.

Kittaka, H. & Tominaga, M. 2017. The molecular and cellular mechanisms of itch and the involvement of TRP channels in the peripheral sensory nervous system and skin. Allergol Int, 66, 22–30.

Lawson, S. N., Mccarthy, P. W. & Prabhakar, E. 1996. Electrophysiological properties of neurones with CGRP-like immunoreactivity in rat dorsal root ganglia. Journal of Comparative Neurology, 365, 355–366.

Lin, S., Liu, X., Jiang, J., Ge, W., Zhang, Y., Li, F., Tao, Q., Liu, S., Li, M. & Chen, H. 2024. The involvement of keratinocytes in pruritus of chronic inflammatory dermatosis. Experimental Dermatology, 33, e15142.

Luostarinen, S., Hamalainen, M. & Moilanen, E. 2021. Transient Receptor Potential Ankyrin 1 (TRPA1)-An Inflammation-Induced Factor in Human HaCaT Keratinocytes. Int J Mol Sci, 22.

Mahmoud, R. H., Brooks, S. G. & Yosipovitch, G. 2024. Current and emerging drugs for the treatment of pruritus: an update of the literature. Expert Opinion on Pharmacotherapy, 25, 655–672.

Mann, C., Dreher, M., Weess, H. G. & Staubach, P. 2020. Sleep Disturbance in Patients with Urticaria and Atopic Dermatitis: An Underestimated Burden. Acta Derm Venereol, 100, adv00073.

Mcdermott, L. A., Weir, G. A., Themistocleous, A. C., Segerdahl, A. R., Blesneac, I., Baskozos, G., Clark, A. J., Millar, V., Peck, L. J., Ebner, D., Tracey, I., Serra, J. & Bennett, D. L. 2019. Defining the Functional Role of NaV1.7 in Human Nociception. Neuron, 101, 905–919 e8.

Meents, J. E., Bressan, E., Sontag, S., Foerster, A., Hautvast, P., Rosseler, C., Hampl, M., Schuler, H., Goetzke, R., Le, T. K. C., Kleggetveit, I. P., Le Cann, K., Kerth, C., Rush, A. M., Rogers, M., Kohl, Z., Schmelz, M., Wagner, W., Jorum, E., Namer, B., Winner, B., Zenke, M. & Lampert, A. 2019. The role of Nav1.7 in human nociceptors: insights from human induced pluripotent stem cell-derived sensory neurons of erythromelalgia patients. Pain, 160, 1327–1341.

Meng, J., Moriyama, M., Feld, M., Buddenkotte, J., Buhl, T., Szollosi, A., Zhang, J., Miller, P., Ghetti, A., Fischer, M., Reeh, P. W., Shan, C., Wang, J. & Steinhoff, M. 2018. New mechanism underlying IL-31-induced atopic dermatitis. J Allergy Clin Immunol, 141, 1677–1689 e8.

MIEßner, H., Seidel, J. & Smith, E. S. J. 2022. In vitro models for investigating itch. Frontiers in Molecular Neuroscience, 15.

Muller, Q., Beaudet, M.-J., De Serres-Bérard, T., Bellenfant, S., Flacher, V. & Berthod, F. 2018. Development of an innervated tissue-engineered skin with human sensory neurons and Schwann cells differentiated from iPS cells. Acta Biomaterialia, 82, 93–101.

Odawara, A., Shibata, M., Ishibashi, Y., Nagafuku, N., Matsuda, N. & Suzuki, I. 2022. In Vitro Pain Assay Using Human iPSC-Derived Sensory Neurons and Microelectrode Array. Toxicological Sciences, 188, 131–141.

Pereira, U., Boulais, N., Lebonvallet, N., Lefeuvre, L., Gougerot, A. & Misery, L. 2010. Development of an in vitro coculture of primary sensitive pig neurons and keratinocytes for the study of cutaneous neurogenic inflammation. Exp Dermatol, 19, 931–5.

Ring, J. 2021. Itch – The major symptom of skin disease and yet still enigmatic. Journal of the European Academy of Dermatology and Venereology, 35, 780–780.

Roggenkamp, D., Falkner, S., Stäb, F., Petersen, M., Schmelz, M. & Neufang, G. 2012. Atopic Keratinocytes Induce Increased Neurite Outgrowth in a Coculture Model of Porcine Dorsal Root Ganglia Neurons and Human Skin Cells. Journal of Investigative Dermatology, 132, 1892–1900.

Roggenkamp, D., Kopnick, S., Stab, F., Wenck, H., Schmelz, M. & Neufang, G. 2013. Epidermal nerve fibers modulate keratinocyte growth via neuropeptide signaling in an innervated skin model. J Invest Dermatol, 133, 1620–8.

Schwartzentruber, J., Foskolou, S., Kilpinen, H., Rodrigues, J., Alasoo, K., Knights, A. J., Patel, M., Goncalves, A., Ferreira, R., Benn, C. L., Wilbrey, A., Bictash, M., Impey, E., Cao, L., Lainez, S., Loucif, A. J., Whiting, P. J., Gutteridge, A., Gaffney, D. J. & Consortium, H. 2018. Molecular and functional variation in iPSC-derived sensory neurons. Nat Genet, 50, 54–61.

Silverberg, J. I., Gelfand, J. M., Margolis, D. J., Boguniewicz, M., Fonacier, L., Grayson, M. H., Simpson, E. L., Ong, P. Y. & Chiesa Fuxench, Z. C. 2018. Patient burden and quality of life in atopic dermatitis in US adults: A population-based cross-sectional study. Ann Allergy Asthma Immunol, 121, 340–347.

Silverberg, J. I., Pinter, A., Alavi, A., Lynde, C., Bouaziz, J. D., Wollenberg, A., Murrell, D. F., Alpizar, S., Laquer, V., Chaouche, K., Ahmad, F., Armstrong, J. M. & Piketty, C. 2021. Nemolizumab is associated with a rapid improvement in atopic dermatitis signs and symptoms: subpopulation (EASI ≥ 16) analysis of randomized phase 2B study. Journal of the European Academy of Dermatology and Venereology, 35, 1562–1568.

Smith, E. S. J., Burton, M. D., Heegaard, A.-M. & Stucky, C. L. 2025. Not Just Neurons: Pain Is Orchestrated in Partnership with Many Non-neuronal Cells. The Journal of Neuroscience, 45, e1309252025.

Ständer, S. & Schmelz, M. 2024. Skin Innervation. Journal of Investigative Dermatology, 144, 1716–1723.

Talagas, M., Lebonvallet, N., Berthod, F. & Misery, L. 2020. Lifting the veil on the keratinocyte contribution to cutaneous nociception. Protein Cell, 11, 239–250.

Tominaga, M. & Takamori, K. 2014. Itch and nerve fibers with special reference to atopic dermatitis: therapeutic implications. J Dermatol, 41, 205–12.

Tsantoulas, C., Farmer, C., Machado, P., Baba, K., Mcmahon, S. B. & Raouf, R. 2013. Probing functional properties of nociceptive axons using a microfluidic culture system. PLoS One, 8, e80722.

Ulmann, L., Rodeau, J.-L., Danoux, L., Contet-Audonneau, J.-L., Pauly, G. & Schlichter, R. 2007. Trophic effects of keratinocytes on the axonal development of sensory neurons in a coculture model. European Journal of Neuroscience, 26, 113–125.

Weisshaar, E. 2021. Itch: A Global Problem? Front Med (Lausanne), 8, 665575.

Wilson, S. R., Nelson, A. M., Batia, L., Morita, T., Estandian, D., Owens, D. M., Lumpkin, E. A. & Bautista, D. M. 2013a. The ion channel TRPA1 is required for chronic itch. J Neurosci, 33, 9283–94.

Wilson, S. R., The, L., Batia, L. M., Beattie, K., Katibah, G. E., Mcclain, S. P., Pellegrino, M., Estandian, D. M. & Bautista, D. M. 2013b. The epithelial cell-derived atopic dermatitis cytokine TSLP activates neurons to induce itch. Cell, 155, 285–95.

Xu, X., Yu, C., Xu, L. & Xu, J. 2022. Emerging roles of keratinocytes in nociceptive transduction and regulation. Frontiers in Molecular Neuroscience, 15.

Yosipovitch, G., Misery, L., Proksch, E., Metz, M., Stander, S. & Schmelz, M. 2019. Skin Barrier Damage and Itch: Review of Mechanisms, Topical Management and Future Directions. Acta Derm Venereol, 99, 1201–1209.

